# Comparative transcriptomics reveal conserved modules of plant defence against different pathogens in Strawberry

**DOI:** 10.1101/2020.06.07.138420

**Authors:** Raghuram Badmi, Arsheed Hussain Sheikh

## Abstract

Strawberry (*Fragaria* ×*ananassa)* is an economically important high-value crop that is susceptible to three most devastating pathogens with different lifestyles – a necrotrophic fungus *Botrytis cinerea* causing grey mold, a hemibiotrophic oomycete *Phytophthora cactorum* causing crown/root rot, and a biotrophic fungus *Podosphaera aphanis* causing powdery mildew. Studies on individual plant-pathogen interactions are only sufficient for developing disease resistant strawberry varieties to a particular pathogen. However, each of these pathogens have the potential to co-infect strawberry at a given point of time. Therefore, understanding how these pathogens manipulate strawberry’s defences and how it responds to these pathogens is essential for developing broad-spectrum disease resistant varieties. Here, in the diploid model *Fragaria vesca*, we performed comparative transcriptome analysis between each of these pathogen infections to identify 501 <u>Co</u>mmon <u>Re</u>sponsive (CoRe) genes targeted against these pathogens. Furthermore, about 80% of these CoRe set are upregulated upon infection by all three pathogens indicating a similar transcriptional response of *F. vesca* independent of pathogen’s lifestyle. These upregulated CoRe set include genes from well-known defence responsive pathways such as calcium and MAP kinase signalling, WRKY transcription factors, pathogenesis-related allergen genes and hormone and terpene biosynthetic genes. These novel insights into *F. vesca*’s defences might serve as a basis for engineering plants with broad spectrum resistance.

## Introduction

At a given time, plants are constantly targeted by multiple biotic stresses like viruses, bacteria, fungi and oomycetes. There exists an evolutionary arms race between plant’s defences and pathogen’s counter defence mechanisms in determining the final outcome of the infection. Tipping the balance in favour of the pathogen results in perturbation of plant’s physiological processes leading to disease spread and eventual plant death. Plant diseases affect crop health and yield amounting to severe loss to the agriculture sector worldwide. Each plant species may be vulnerable to attack by several pathogens with different lifestyles causing variety of diseases. Plants can mount unique defences against a particular pathogen, however, there are common features where a robust immunity response is generated to fight many related pathogens at a single time. For example, a particularly high value crop strawberry is susceptible to three pathogens with different lifestyles:

i. *Botrytis cinerea* – a necrotrophic fungi and a generalist pathogen with a broad host range which is capable of infecting more than 1000 plant species (Petrasch et al., 2019)
ii. *Phytophthora cactorum* – a hemibiotroph and an oomycete that causes crown/root rot in strawberry (Toljamo et al., 2016). Other species of *Phytophthora* infect important crop species such as potatoes causing the devastating late blight disease (Yang et al., 2017).
iii. *Podosphaera aphanis* – a biotrophic fungus that causes powdery mildew in strawberry, apple, grapevine, wheat and barley and other important crops (Kiss et al., 2004).

Given that strawberry has multiple natural pathogens that damage the crop, no study has yet been conducted to integrate the information from different pathogen responses to derive new knowledge that provide insights into the conserved defence mechanisms in strawberry. Each of these pathogens are individually capable of causing devastating damage to the crops. Understanding the mechanisms of pathogen invasion and spread is crucial for designing appropriate agronomic solutions. However, increasing resistance against one of these pathogens, for example grey mold, might result in decreasing resistance (or increasing susceptibility) against powdery mildew. This is because, it is well known that jasmonic acid (JA) -mediated resistance against necrotrophs (*B. cinerea*) is antagonistic to salicylic acid (SA) -mediated defences against biotrophs (*P. aphanis*) (Thaler et al., 2012). It is also argued that plants tightly control JA- and SA-mediated defences in order to prevent unfavourable molecular interactions in their attempt to resist further attack from pathogens (Spoel et al., 2007). This means that uncontrolled activation of JA-mediated defences would provide resistance against *B. cinerea* infection but also might increase susceptibility against powdery mildew infection by suppressing SA-mediated defences. Therefore, achieving resistance against broad spectrum of pathogens require deeper knowledge about plant’s responses towards these pathogens. Furthermore, comparing the plant’s defence responses against these pathogens will help in deriving shared consensus from a plant’s perspective. More importantly, knowledge from such integrative studies hold the potential for developing plants that display broad spectrum resistance against multiple pathogens.

Transcriptomic responses are one of the most widely studied aspects in plant-pathogen interactions as they offer significant insights into the plant’s immunity. Increasing interest to study transcriptomics of crop-pathogen responses and decreasing costs of transcriptomic studies have resulted in accumulation of wealth of data that is ready to be analysed. It is usually tempting to compare if a certain gene is similarly up- or downregulated upon infection by a different pathogen. Gene ‘X’ upregulated in both *F. vesca* - *B. cinerea* (*FvBc*) pair and *F. vesca* – *P. cactorum* (*FvPc*) pair might mean that the gene’s response against both these pathogen attacks is same. However, if an upregulated gene ‘Y’ in *FvBc* pair is downregulated in *FvPc* pair, it would mean that the gene’s response is different or rather opposite to both these pathogens. Identifying these similarities and differences between different *F. vesca* – pathogen (*FvPathogen*) pairs will provide fundamental insights about *F. vesca*’s defences against broad spectrum of pathogens. However, different gene IDs used in different studies and usage of different transcriptome template for mapping and analysis limits such comparative analysis of results from different pathogen infections. For example, studies mapped to the older transcriptome (Shulaev et al., 2011) used gene IDs in the format geneXXXXX. However, the latest *F. vesca* transcriptome (Edger et al., 2018) uses nomenclature in the format FvH4_XgXXXXX and do not show complete one-to-one match with the previous gene sequences and/or annotations. (Re)Mapping the RNA-seq results to a single transcriptome and using the common pipeline for mapping will provide results that can be reliably compared across other studies and between different pathogen infections. In this article, we mapped the available transcriptomic datasets of *FvPc* (Toljamo et al., 2016) and *F. vesca* – *P. aphanis* (*FvPa*) (Jambagi et al., 2015) pairs to the recent transcriptome version used for mapping the *FvBc* pair (Badmi et al., 2019). The results obtained in this study allows for easy and direct comparison of expression values for the three most important *FvPathogen* pairs. Furthermore, by highlighting the set of genes common between infections, this study highlights the conserved defence mechanisms in *F. vesca* that are responsive to the pathogen infection independent of pathogen lifestyle.

## Materials and Methods

### RNA-seq data retrieval and processing

The SRA files of raw RNA-seq reads were downloaded using ‘fastq-dump’ command in windows command line terminal. Quality control, adapter trimming and quality filtering were performed using fastp program (Chen et al., 2018). The resulting reads were then aligned to the latest *F. vesca* transcriptome (Edger et al., 2018) (ftp://ftp.bioinfo.wsu.edu/species/Fragaria_vesca/Fvesca-genome.v4.0.a1) using rsem aligner with bowtie2 option (Li et al., 2011). From the gene results files, Fragments per kilobase of exon per million reads mapped (FPKM) values were extracted from and analysed for differentially expressed genes using EdgeR (McCarthy et al., 2012; Robinson et al., 2009). The ‘trimmed mean of M values’ (TMM) method was used for normalization and Benjamini and Hochberg approach was used to adjust the p-values (Benjamini and Hochberg., 1995).

### Analysing differentially expressed genes

The differentially expressed genes from four *F. vesca* – pathogen interactions (*FvBc, FvPc* and *FvPa*) were compiled in a single spreadsheet with the same *F. vesca*_4.0 IDs and filtered to retain the expression values that have >1.5 log2 fold change values. For the *FvBc* and *FvPc* pair, two filters were used: the p-adjusted values < 0.05 and > 1.5 log2 fold change were retained. These sets of differentially expressed genes were used to generate venn diagrams using InteractiVenn online tool (Heberle et al., 2015). The list of genes intersecting between the three pairs *FvBc, FvPc* and *FvPa* were retrieved from the venn diagram by clicking on the intersecting area. This list was named as CoRe set and was used for further analysis. The 3D scatter plot to visualize scatter of log2FC values between these three sets was plotted using Matplotlib tool (https://matplotlib.org/) in Python 3.6.

## Results

### Transcriptome analysis of *Fragaria vesca* – pathogen interactions

The RNA-seq data from *FvBc* interactions were mapped to the latest *F. vesca*_4.0.a1 transcriptome (Badmi et al., 2019), whereas *FvPc* (Toljamo et al., 2016) and *FvPa* (Jambagi et al., 2015) were mapped to older version (Shulaev et al., 2011). To be able to compare these three *FvPathogen* interactions, the RNA-seq reads from *FvPc* and *FvPa* were mapped to the same *F. vesca*_4.0.a1 reference transcriptome. The data for *FvBc* pair was used from Badmi et al (2019). For *FvPa* pair, only the dataset representing 1 day after infection by *P. aphanis* and using *F. vesca* Hawaii-4 as a host was considered for analysis, since this was appropriate for comparing with other chosen datasets. Table 1 lists the accession numbers of these datasets and corresponding sample names. A total of 28,588 transcripts were mapped for each dataset. The complete list of log2FC values for each pathogen infection with a single *F. vesca* gene IDs are presented in Supplementary Table 1. The differentially expressed transcripts were filtered based on the criteria of log2FC > 1.5 and p < 0.05 (Supplementary Table 2). The retained log2FC values were used for further analysis. Based on the above criteria, 10.17% of the *F. vesca* transcriptome was differentially expressed including 1535 upregulated and 1373 downregulated genes upon *B. cinerea* infection. *P. cactorum* infection resulted in differentially expression of 11.41% of the *F. vesca* transcriptome which included 1605 upregulated genes and 1659 downregulated genes. Infection by *P. aphanis* differentially expressed 12.89% of the *F. vesca* transcriptome consisting of 3024 up- and 660 downregulated genes (Table 1, Fig 1).

**Table 1:** Details of the genotype, pathogen, samples and their SRA accession numbers used for mapping the RNA-seq reads to the *F. vesca* transcriptome and their corresponding number of differentially expressed genes.

| *F. vesca*<br>genotype | Pathogen | RNA-seq<br>sample | NCBI SRA<br>accession<br>numbers | Upregulated<br>log2FC ><br>1.5 | Downregulated<br>log2FC > 1.5 | P<br>adjusted | Reference |
| --- | --- | --- | --- | --- | --- | --- | --- |
| Hawaii 4 | *Botrytis*<br>*cinerea* | n.a. | n.a. | 1535 | 1373 | < 0.05 | (Badmi et<br>al., 2019) |
| Hawaii 4 | *Phytophthora*<br>*cactorum* | Control 1 | SRR3743193 | 1605 | 1659 | < 0.05 | (Toljamo et<br>al., 2016) |
|  |  | Control 2 | SRR3743194 |  |  |  |  |
|  |  | Control 3 | SRR3743195 |  |  |  |  |
|  |  | *P. cactorum*<br>inoculated 1 | SRR3743196 |  |  |  |  |
|  |  | *P. cactorum*<br>inoculated 2 | SRR3743197 |  |  |  |  |
|  |  | *P. cactorum*<br>inoculated 3 | SRR3743198 |  |  |  |  |
| Hawaii 4 | *Podosphaera*<br>*aphanis* | Control | ERR419272 | 3024 | 660 | n.a. | (Jambagi &<br>Dunwell.,<br>2015) |
|  |  | *P. aphanis*<br>inoculated | ERR419273 |  |  |  |  |

**Figure 1:**
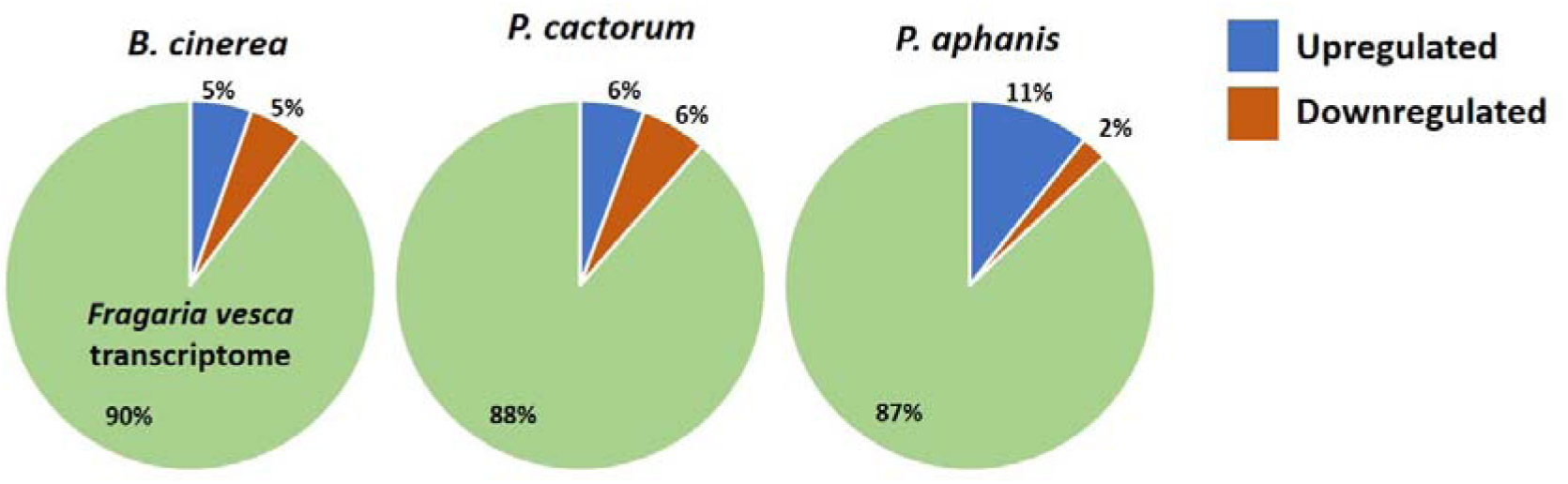
Percentage of *Fragaria vesca* transcriptome dysregulated by pathogens. Pie charts representing the percentage of up- and down-regulated genes as a percentage of *F. vesca* transcriptome. Blue cone represents the upregulated percentage, orange cone represents downregulated percentage and pale green part represents the un-regulated transcriptome.

### Common Responsive (CoRe) genes of *Fragaria vesca* against pathogens

To identify commonly regulated genes between three *Fvpathogen* pairs, venn diagram was generated using the filtered log2FC gene list of *FvBc, FvPc* and *FvPa* (Supplementary table 2) (Fig. 2). A total of 2908 genes from *B. cinerea* infection, 3684 genes from *P. aphanis* infection and 3264 genes from *P. cactorum* infection were used for generating venn diagram. Interestingly, the intersection of all three pathogen infections resulted in 501 genes (Fig. 2) representing the CoRe set that respond to all three pathogen infections.

**Figure 2:**
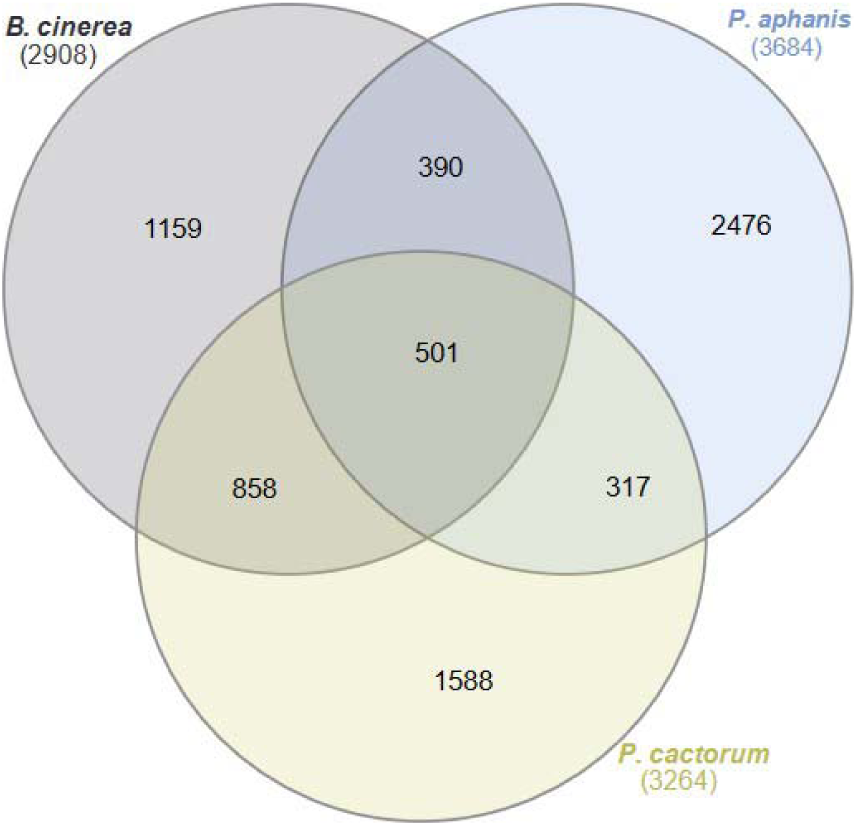
Venn diagram of the dysregulated genes in the three *F. vesca* – pathogen pairs showing the number of specifically and commonly regulated genes between each pair. The numbers represent the number of genes included in respective sets. The central part representing 501 genes represents the Commonly Regulated genes or CoRe set.

The CoRe genes were retrieved from InteractiVenn by clicking on the intersecting area between the three pathogen infections (Supplementary table 3). The CoRe list represents genes that are common between all three *FvPathogen* pairs, including both upregulated and downregulated genes. As expected, the CoRe genes consisted of well-known defence responsive genes from secondary metabolite pathways, receptor kinases, WRKY transcription factors and hormone pathways. Among the CoRe genes, 429 genes were upregulated and 72 genes were downregulated in *FvBc* sample (Fig. 3a). In *FvPc* sample, 425 genes were upregulated and 76 genes were downregulated. And in *FvPa* sample, 457 genes were upregulated and 44 genes were downregulated. Clearly, all three pathogen infections had higher number of upregulated genes in the CoRe list. To visualise the overview of the gene expression patterns in all three samples, a 3D scatter plot was generated using MatPlotLib tool in Python 3 (Fig. 3b). The scatter plot represents 501 genes with each dot representing a single gene and its position in the plot is determined by the expression level of that gene in the three *FvPathogen* interactions. As expected from the higher number of upregulated genes, most of the genes clustered in the area representing upregulation in all the three pathogen responses (Fig. 3b). Out of 501 CoRe genes, 400 genes were upregulated in all the three *FvPathogen* pairs (Supplementary Table 3), pointing out that these genes are commonly upregulated independent of pathogen lifestyle. However, only 26 genes in the core set were downregulated in all three pairs and 75 genes displayed either up- or downregulation (mixed regulated).

**Figure 3:**
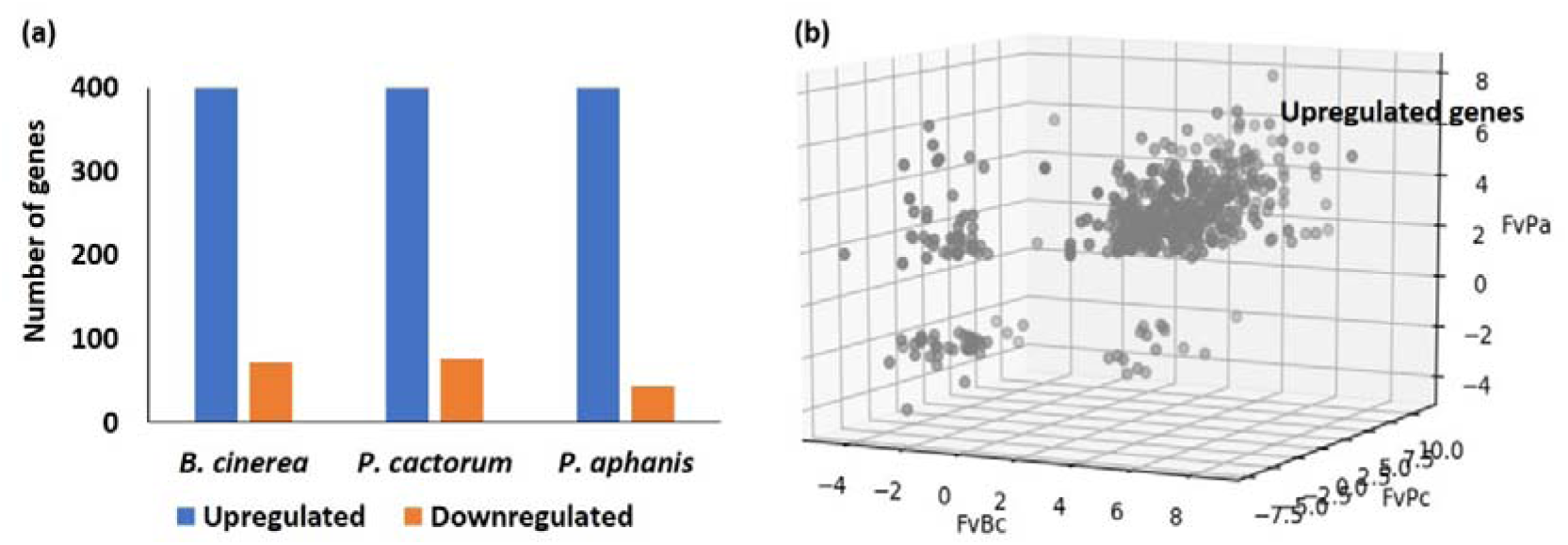
(a) A bar graph representing the number of Commonly Regulated (CoRe) set of genes up- and down-regulated in each *F. vesca* – pathogen pair. (b) A 3D scatter plot representing 501 CoRe genes with each dot representing a single gene. The three *F. vesca* – pathogen pairs, *FvBc, FvPa* and *FvPc* form X-, Y- and Z-axis respectively. Each dot representing each gene is positioned relative to its expression levels in the three *F. vesca* – pathogen pairs.

## Discussion

One of the key characteristics of *F. vesca* is its natural susceptibility to pathogens as diverse as biotrophs, necrotrophs and hemibiotrophs. As these plant-pathogen pairs are of natural origin, studying their interactions will provide insights that are highly relevant for extension to other crop-pathogen interactions. Additionally, this opens opportunities to understand how plants identify and respond to pathogens with different lifestyle in a natural context. The difference in the percentage of transcriptome dysregulation upon each pathogen infection indicates that each pathogen reprograms the host’s transcriptome to a different extent. This probably means that the overall effect of a pathogen on the plant’s transcriptome is different although there might be some overlapping transcripts regulated commonly (CoRe) between these three pathogens. This difference might also be due to the perception of specific pathogen or microbe associated molecular pattern (PAMP or MAMP) and/or the specific effectors secreted by the individual pathogens that determine the nature of downstream differential expression of genes. Identifying the common overlap between these three pathogen infections will illuminate the conserved defence responses of *F. vesca* that function independent of pathogen lifestyle.

The commonly upregulated genes in CoRe set include well known defence responsive genes such as MAPK and Ca2+ signalling, Ca2+ binding, calmodulin proteins, WRKY transcription factors, pathogenesis-related allergen genes, genes involved in defence hormone biosynthesis and terpene biosynthesis. MAPK cascades are one of the first responders to pathogen infection, that transduce signals perceived on the extracellular surfaces to the cytoplasmic and nuclear factors resulting in appropriate pathogen responses (Pitzschke et al., 2009). Upregulation of MAPKKK encoding genes indicates the importance of these genes in signal transduction upon perception of these three pathogens. Additionally, *WRKY* transcription factors, the well-known substrates of MAPK cascades (Eulgem and Somssich., 2007; Popescu et al., 2009; Sheikh et al., 2016), are also upregulated in the core set thus reinforcing the importance of MAPK signalling. MAPK signalling (Raina et al., 2012) and WRKY transcription factors (Frerigmann et al., 2016) are independently shown to be involved in secondary metabolism. Calcium signalling pathways plays vital roles in pathogen responses (Tian et al., 2019), symbiotic associations (Vadassery et al., 2009) and cell-wall biosynthesis (Badmi et al., 2018). The upregulation of genes encoding calcium binding proteins, calmodulin and calmodulin binding IQ-domain (IQD) proteins points to their important roles in defence responses against these three pathogens. Other stress related transcription factors such as *JUNGBRUNNEN* (Thirumalaikumar et al., 2018) and Zinc finger proteins (Maldonado-Bonilla et al., 2013) are also upregulated in the CoRe set. The activated transcription factors further activate the metabolite genes promoting the biosynthesis of hormones and secondary metabolites (Zhou et al., 2016). The upregulated CoRe set also includes genes encoding 12-oxophytodienoate reductase (12-OR2) and 1-aminocyclopropane-1-carboxylate (ACC) oxidase that are involved in the biosynthesis of jasmonic acid and ethylene respectively, the two most important defence hormones (Park et al., 2018; Schaller et al., 2000). Furthermore, the genes encoding (-)-germacrene D synthase (GDS), 3-hydroxy-3-methylglutaryl-coenzyme A (HMG CoA) reductase and beta-amyrin oxidase are also upregulated in the CoRe set that are respectively involved in the biosynthesis of sesquiterpenes (Schmidt et al., 1998), mevalonic acid (Weissenborn et al., 1995) and the triterpene ginsenoside (Han et al., 2013). These observations point out that these specific genes (i) commonly respond to pathogen attacks independent of pathogen lifestyle and (ii) are probably involved in mediating signal transduction that converges on leading to biosynthesis of defence related secondary metabolites. This prompts to envisage a working signal transduction pathway that is activated in response to broad spectrum pathogen infections (Fig. 4). Literature evidence supports the upregulation of these genes to the biological functions observed in defence responses against pathogens. More research on this front is required to address the outstanding questions such as (i) do the upregulated CoRe genes function as part of a gene regulatory network that is important in *F. vesca* disease resistance? (ii) does the transcriptional upregulation reflect increased accumulation of defence related secondary metabolites in response to pathogen attack?

**Figure 4:**
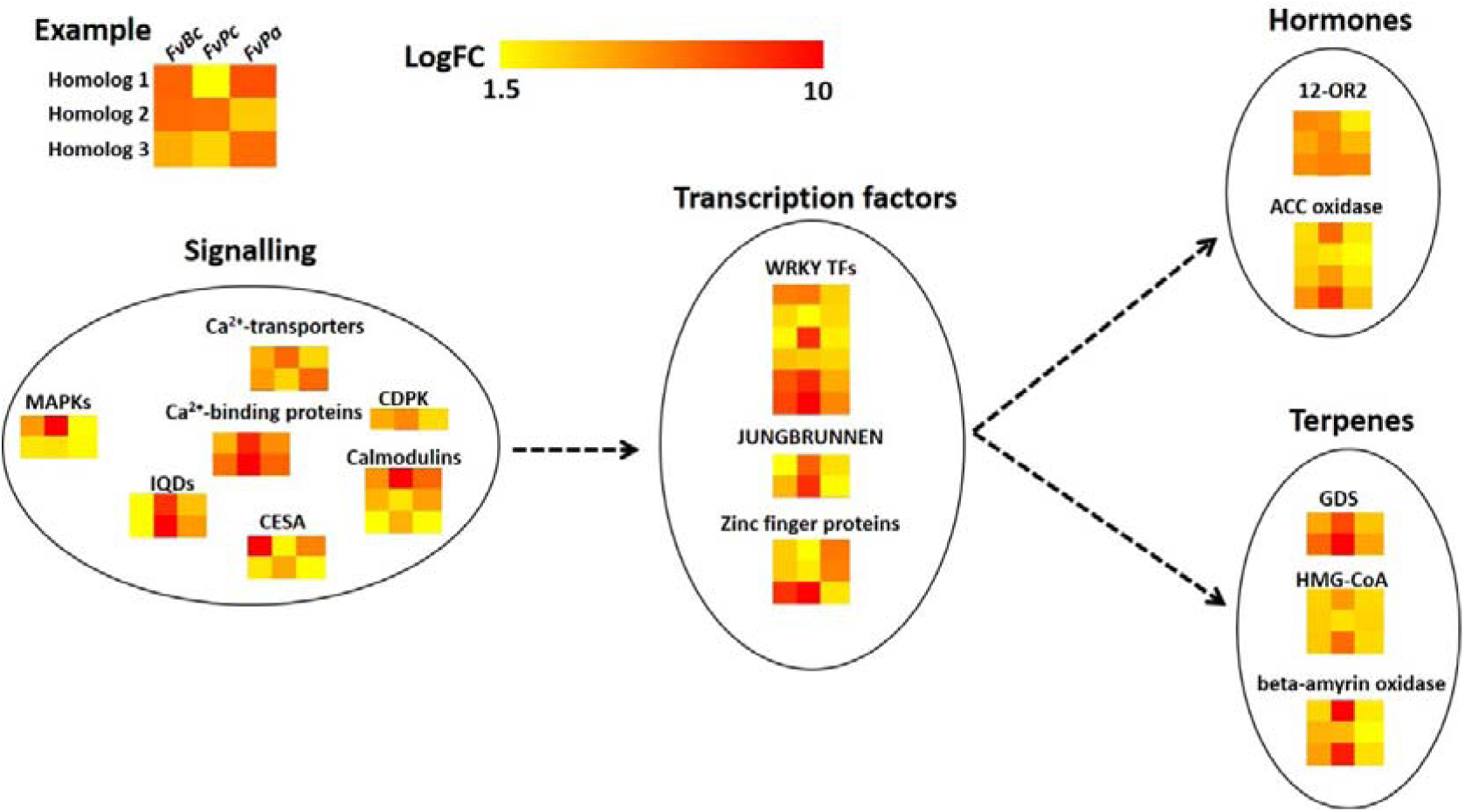
A probable heatmap signalling pathway that reflects the transcriptional upregulation of genes in the three *F. vesca* – pathogen pairs. Each row of the heatmap represents a single gene and each column corresponds to the respective *F. vesca* – pathogen pair as shown in the example.

## Author Contributions

RB conceived the idea, retrieved and analysed the data. RB and AHS wrote the manuscript

